# Bioluminescence and fluorescence of three sea pens in the north-west Mediterranean sea

**DOI:** 10.1101/2020.12.08.416396

**Authors:** Warren R Francis, Anaïs Sire de Vilar

## Abstract

Bioluminescence of Mediterranean sea pens has been known for a long time, but basic parameters such as the emission spectra are unknown. Here we examined bioluminescence in three species of Pennatulacea, *Pennatula rubra, Pteroeides griseum*, and *Veretillum cynomorium*. Following dark adaptation, all three species could easily be stimulated to produce green light. All species were also fluorescent, with bioluminescence being produced at the same sites as the fluorescence. The shape of the fluorescence spectra indicates the presence of a GFP closely associated with light production, as seen in *Renilla*. Our videos show that light proceeds as waves along the colony from the point of stimulation for all three species, as observed in many other octocorals. Features of their bioluminescence are strongly suggestive of a “burglar alarm” function.

## Introduction

Bioluminescence is the production of light by living organisms, and is extremely common in the marine environment [Haddock et al., 2010, Martini et al., 2019]. Within the phylum Cnidaria, biolumiescence is widely observed among the Medusazoa (true jellyfish and kin), but also among the Octocorallia, and especially the Pennatulacea (sea pens).

Bioluminescence of Pennatulacea in the Mediterranean has been known for centuries (reviewed by [Harvey, 1957]). The first detailed observations on several Mediterranean sea pens were made in the 19th century, by [Panceri, 1872a]. This work had examined *Pennatula rubra* (Ellis 1761), *Pennatula phosphorea* (Linnaeus 1758), *Funiculina quadrangularis* (Pallas 1766), and *Pteroeides griseum* (Bohadsch 1761), and had described the rapid propagation of light along the colony following stimulation [Panceri, 1872a]. It was not known at this time if this was under the control of a nervous system [Panceri, 1872a], or if a nervous system was even present in Cnidarians.

Later observations on the bioluminescence of several of the same species were done by Titschack [Titschack, 1964, Titschack, 1966] in the 1960s. Working on *P. rubra, P. griseum* and *Veretillum cynomorium* (Pallas 1766), it was noted that the fluorescence and bioluminescence originated from the same cells, suggesting a close connection between the green bioluminescence and the fluorescence. Furthermore, like all other sea pens, it was later demonstrated that the luminescence is under nervous control [Bilbaut, 1975a,Bilbaut, 1975b], arguing that the light is not merely a metabolic by-product.

Biochemically, the bioluminescence of Pennatulacea has been best studied in the sea pansy *Renilla reniformis*. This involves two key components, the luciferin, coelenterazine, and the enzyme *Renilla* luciferase. These two components are necessary and sufficient for light production, though in the natural system, two additional proteins are found. One is a calcium-activated luciferin-binding protein (LBP) for coelenterazine [Anderson et al., 1974], and the other is a green fluorescent protein (GFP) that interacts closely with the luciferase [Ward and Cormier, 1978] to modify the color of the emitted light by resonant energy transfer. It was found that extracts from many sea pens can create light when combined with extracts from other species [Cormier et al., 1973, Wampler et al., 1973], suggesting that all sea pens share the same biochemical system. Additionally, all of the species examined by [Wampler et al., 1973] produced green light due to interaction of the luciferase to a GFP.

Because much of the early work on sea pen bioluminescence was descriptive, and the photographs taken at the time are difficult to interpret, there is a need for an updated examination and interpretation of the role of bioluminescence for these animals. Several Mediterranean sea pens (*Pennatula rubra, Pennatula phosphorea*, and *Pteroeides griseum*) are currently listed as “vulnerable” by the IUCN [del Mar Otero et al., 2017]. Our understanding of the biology and ecology of sea pens is more important now than ever before. Thus, here we examined the fluorescence and bioluminescence of three sea pen species in the north-west Mediterranean. We have observed that all three species produce green light when disturbed, which could serve a range of functions during encounters with different predators.

## Methods

### Specimens

Specimens of *Pennatula rubra, Pteroeides griseum*, and *Veretillum cynomorium* were collected around the Bay of Banyuls in early October 2020. All three species studied are colonial organisms that live in sandy or muddy habitats. The colony consists of a peduncle that buries itself in the substrate and a rachis containing multiple types of zooids, depending on the species.

A total of 8 specimens of *Veretillum cynomorium* were collected by divers in the bay in the morning Oct-2, at a depth of 25m, appx 42.487N 3.145E. One specimen was also collected by dredge, along with the other sea pens. A total of 13 specimens of *Pennatula rubra* and 28 specimens of *Pteroeides griseum* were collected by dredge on Oct-8 at a depth of 55-60m, 42.49N 3.16E.

All experimental work was carried out at the Laboratoire d’Arago of Banyuls-sur-mer. Specimens were maintained in 60L aquarium tanks with a constant flow of filtered seawater (200 micron filter) at ambient temperature, typically 17-20 degrees.

### Fluorescence spectra

The *in vivo* fluorescence spectra were measured using an Ocean Optics HR4000CG-UV-NIR spectrophotometer, with attached fiber optic. Specimens were illuminated with a blue diving flashlight (peak wavelength: 450nm), and the spectrum was captured through a yellow filter. We used the program brizzy (https://github.com/conchoecia/brizzy) modified from the Ocean Optics SeaBreeze API. As the sensitivity of the spectrophotometer was relatively low, long exposure (10-15s) was necessary to capture the spectra. Spectra were smoothed using the `smooth.spline` function with 25 degrees of freedom, implemented in R.

### Photos and video

Fluorescence and brightfield photos were taken with a Nikon D3200 camera. A Tiffen Yellow 12 lens filter was used for the fluorescence images. Laboratory images and video of bioluminescence were taken with a Canon Rebel SL2, equipped with a Tamron SP AF17-50mm F/2.8 DiII LD Aspherical (IF) Lens. For all photos, settings were f2.8, ISO 6400, and either a 2s or 3.2s exposure. For video, settings were 30fps and ISO 25400. Experiments were carried out in a dark room, where all animals were dark-adapted for at least 30 minutes. The animals were removed from the aquarium and placed into a glass or plastic container for imaging. For photos, bioluminescence was stimulated either by touching the animal with fingers, a glass pipette, or by addition of 7% KCl solution. For the supplemental video, addition of appx. 0.5mL of KCl solution was used to initiate bioluminescence.

## Results

### General observations of the luminescence and fluorescence

The bioluminescence and fluorescence was examined in three species of pennatulacea: *Pennatula rubra* (Figure 1), *Pteroeides griseum* (Figure 2), and *Veretillum cynomorium* (Figure 3). All three species were bioluminescent, emitting green light that was easily visible to dark-adapted eyes. Several observations were common to the three species.

**Figure 1:**
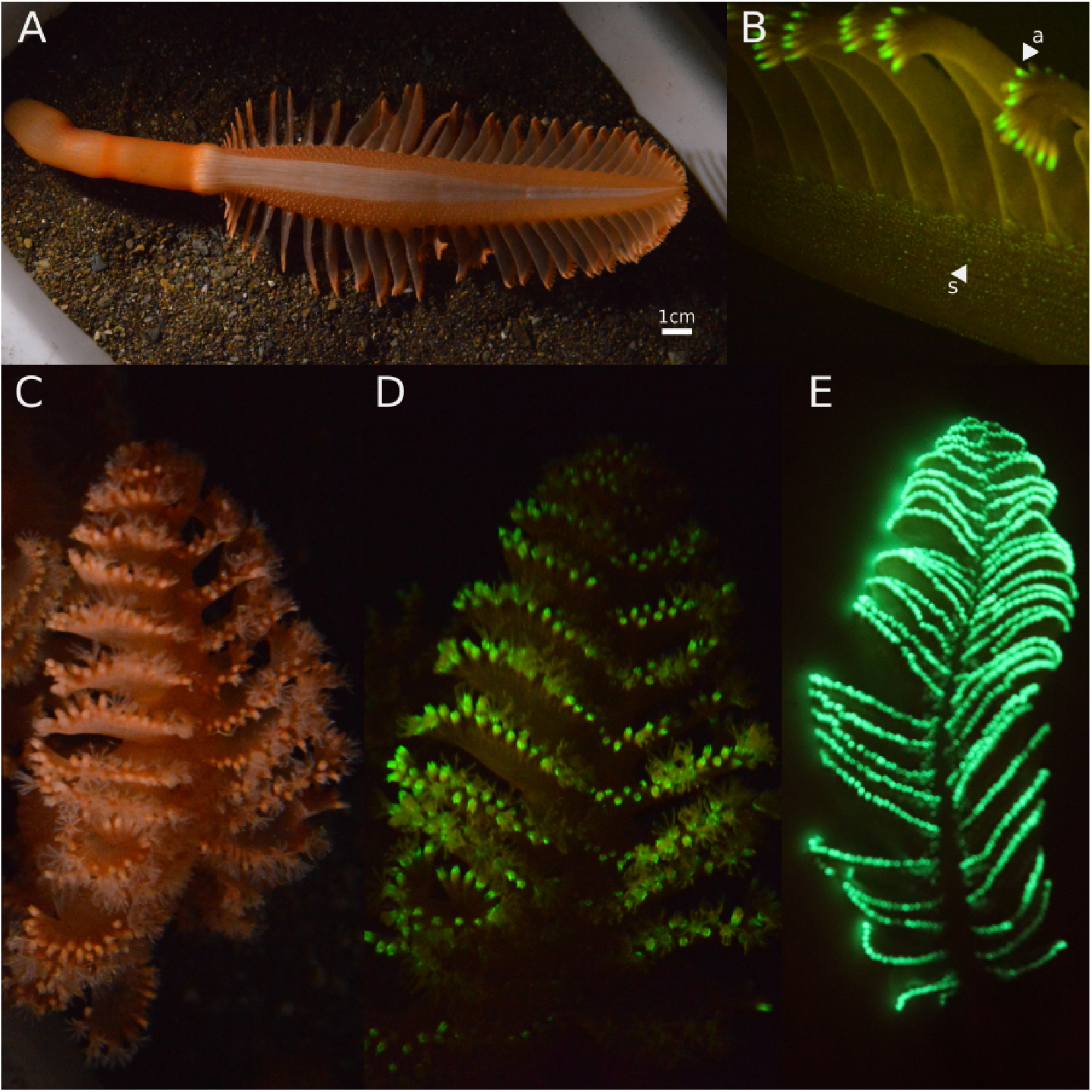
Photos of *Pennatula rubra* (**A**) Posterior view of a normal specimen when inflated, with the characteristic white stripe along the rachis, and siphonozooids visible as small bumps along the rachis. (**B**) Close up view of the fluorescence of the: s-siphonozooids, and a-autozooids, illuminated with blue light using a yellow filter. (**C-D**) Anterior view of a normal specimen with the autozooid tentacles fully extended, under white light (C) and fluorescence (D). The stalk of the autozooids is brightly fluorescent, however, the tentacles of the autozooids do not appear to be fluorescent. (**E**) 2s exposure showing the anterior view of the bioluminescence of another specimen.

**Figure 2:**
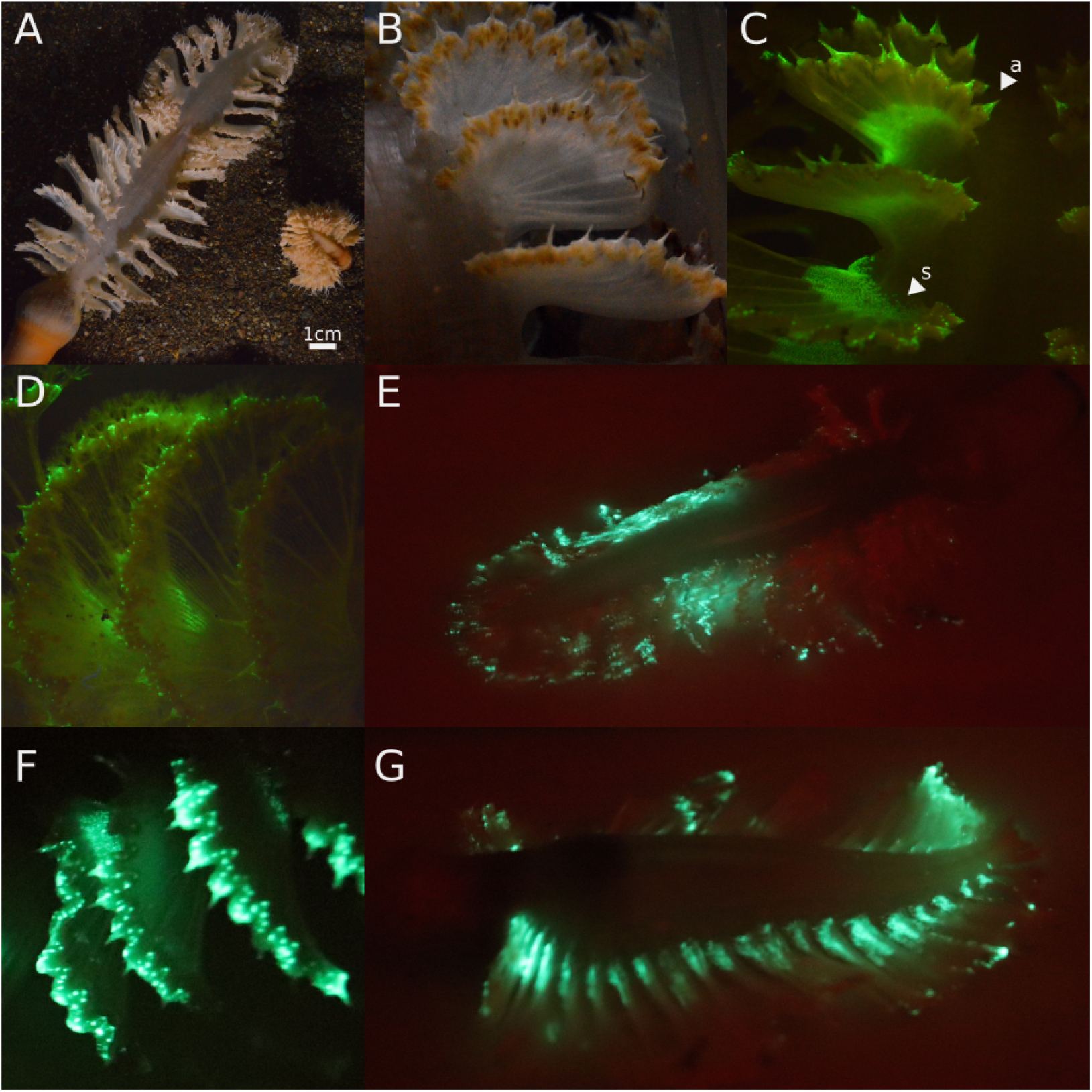
Photos of *Pteroeides griseum* (**A**) Image of two colonies, a “gray” and a “beige” variant, based on the color of the rachis. (**B**) Close up photo of the polyp leaves of, under white light (B), and fluorescence with blue light and a filter (C). The distal autozooids-a concentrated around the spicules and density of siphonozooids-s are both fluorescent. (**D**) Close up image of the polyp leaves when the autozooid tentacles are extended, showing that they are not fluorescent. (**E**) Image of the bioluminescence of another specimen. (**F**) Close up image of the bioluminescence of the autozooids on the polyp leaf. (**G**) Posterior view of the bioluminescence, showing the light from the siphonozooids on the polyp leaves.

**Figure 3:**
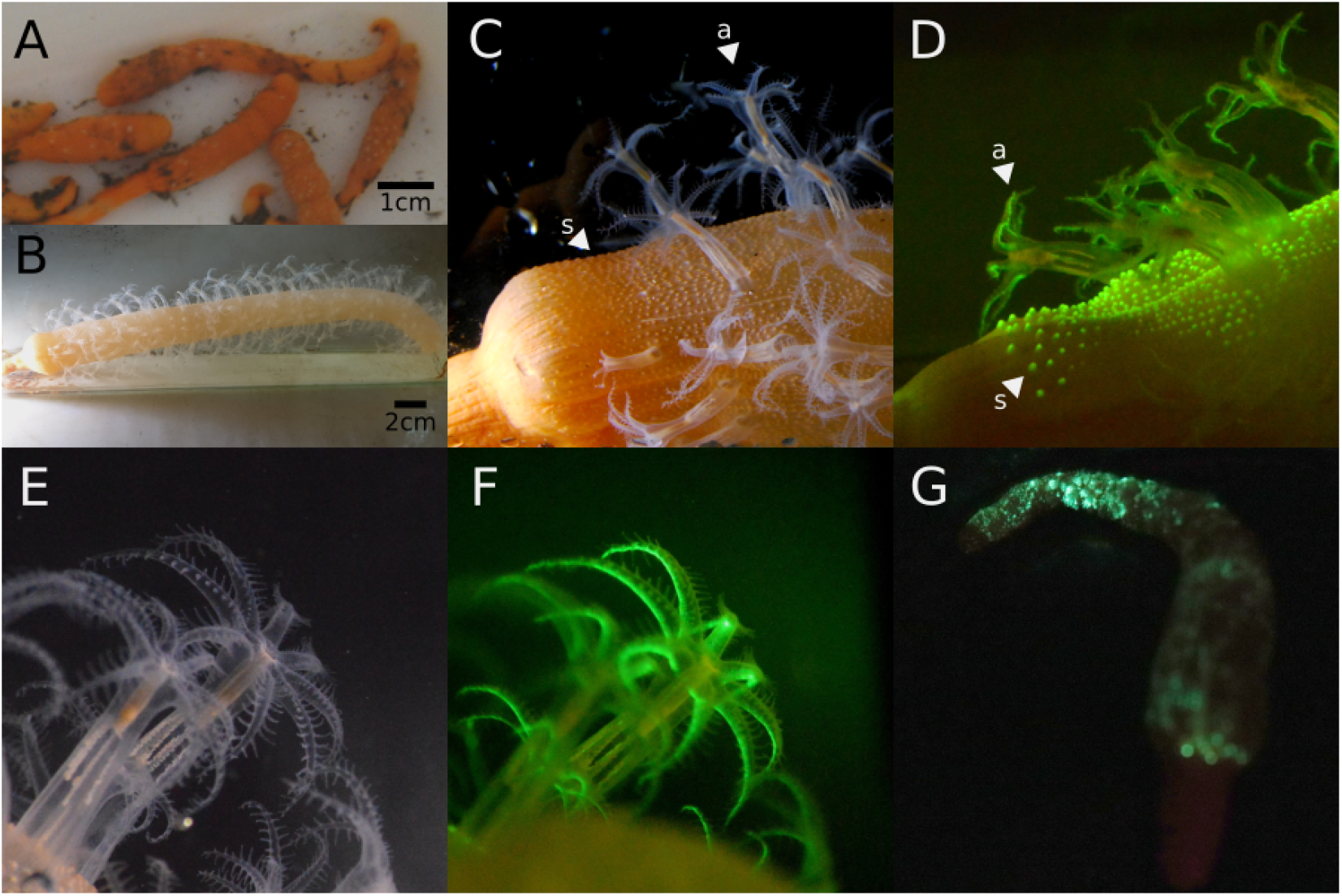
Photos of *Veretillum cynomorium* (**A**) Specimens in fully contracted state, immediately after collection. (**B**) Specimen approximately a day later, in a fully inflated state. (C) Closeup photo of the proximal portion of the rachis, showing the: s-siphonozooiods and a-autozooids, which are also fluorescent (**D**) under blue light with a filter. (**E,F**) Close-up of the autozooids as brightfield, showing that the fluorescence is mostly restricted to the tentacles. (**G**) Bioluminescence of a partially contracted whole specimen, where the peduncle is located at the bottom of the image.

Light emission only occurs in dark-adapted specimens, but animals appeared to be luminous (or could be stimulated) regardless of time of day. That is, if the room was kept dark, specimens were luminous even during daylight hours of the following day. This has been reported in other Pennatulacea species [Davenport and Nicol, 1956].

Fluorescence was dim (i.e. barely visible without a filter), and emission of bioluminescence occurred from the same sites that were fluorescent, confirming the reports by [Titschack, 1964, Titschack, 1966]. The shape of the fluorescence spectra (Figure 4) is indicative of the presence of GFP [Wampler et al., 1973], as seen in many other octocorals [Bessho-Uehara et al., 2020].

**Figure 4:**
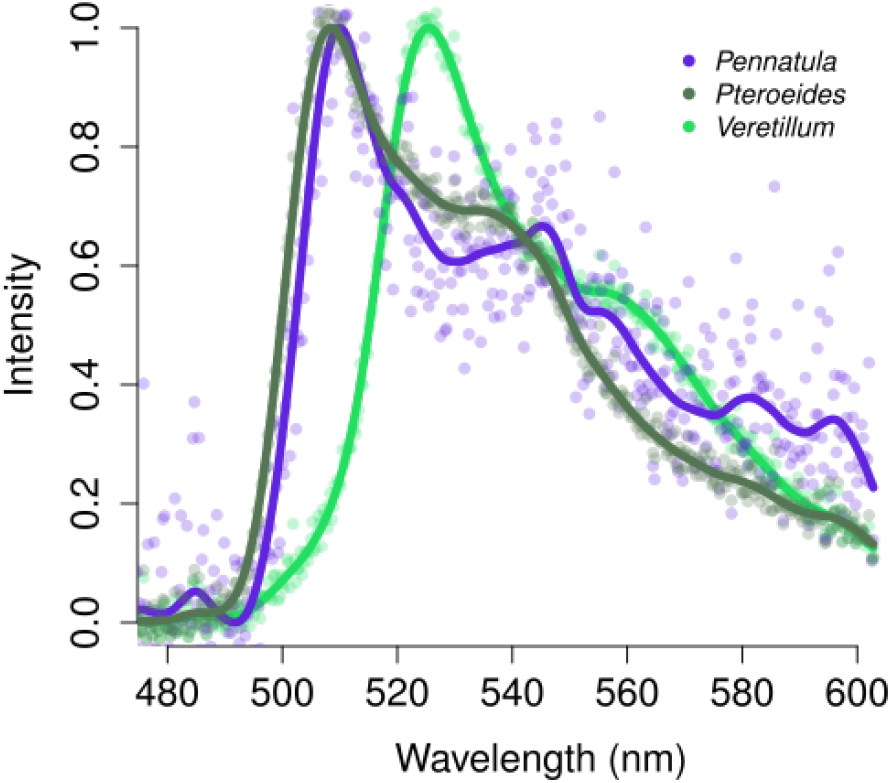
Fluorescence spectra of all three species, showing the characteristic GFP-like shoulder into longer wavelengths. Lines show the smoothed spectra, while the faded points indicate the original capture. Max emission was 509nm for *Pennatula*, 508nm for *Pteroeides*, and 525nm for *Veretillum*.

Light production would begin at the site of stimulation, and waves of light travel outward from the point of stimulation (see Supplemental Video). Light could also be initiated by addition of KCl. When added towards the peduncle or apex, the waves of light passed towards the other end. Drops of KCl applied across the colony (outside of the aquarium) could usually stimulate most of the colony to emit light, but the waves ceased, indicating that there is likely some interference if stimulated at too many sites. Brief flashing routinely occurred even after stimulation has ceased. We had also observed that some manner of colony retraction always occurred following bioluminescence responses, starting with the retraction of the autozooids and then with the total contraction of the rachis.

### Pennatula rubra

From dredging, a total of 13 specimens were collected, of length around 7cm when fully contracted to 26cm when fully extended. This species is in the shape of a feather or fern, and is reddish-orange coloured with a thick central white stripe along the rachis (Figure 1A). All specimens were confirmed to be *Pennatula rubra* on the basis of morphological characteristics reported by [Chimienti et al., 2015] to distinguish them from the related *Pennatula phosphorea*.

Green fluorescence was visible along the autozooids around the edge of the polyp leaves (Figure 1B, 1D), but sparsely along the rachis from the siphonozooids (Figure 1B). Bioluminescence was reliably stimulated by pressing a pipette along or between the polyp leaves. Light initiates from the point of stimulation, but the waves passed very quickly across the entire length of the animal, lighting up along the edge of the polyp leaf, i.e. the autozooids (Figure 1E, Supplemental Video). Directly touching the rachis also initiated light on the siphonozooids, but did not appear to induce the waves of light. All of the mechanical stimuli resulted in rapid contraction, first by withdrawal of the autozooids, then curling up of the whole animal along the rachis, and subsequent deflation. Similar behavior was also observed in the wild [Chimienti et al., 2018]. The curling of the colony produced a shape of a question mark, with all of the autozooids on the polyp leaves pointing outward, and would enable light to be emitted in all directions.

### Pteroeides griseum

From the dredge, 28 specimens of *Pteroeides griseum* were collected, of approximately 6-10cm long when contracted up to 32cm when fully extended (Figure 2A). All specimens were identified as *Pteroeides griseum*, which has been synonymized with *Pteroeides spinosum* [Williams, 1995], though old reports list both species in the Mediterranean. Like *Pennatula*, this species has a general shape of a feather, and bears numerous parallel polyps leaves on each side of the rachis. Unlike *Pennatula*, siphonozooids appear to be absent along the rachis, and are instead concentrated at the base of the polyp leaves (Figure 2B).

Green fluorescence was seen along the tips of the polyp leaf, and among a high density area of siphonozooids on the leaf, but not among the tentacles of the autozooids (Figure 2B-D). Touching the polyp leaf with a pipette produced rapid flashes for a few seconds along the autozooids on the polyp leaf and among density of siphonozooids, while no luminescence was at all visible along the rachis. With repeated or intense stimulation, waves of light extended (relatively) slowly from the point of stimulation (Figure 2E-G, Supplemental Video).

### Veretillum cynomorium

The specimens of *Veretillum* ranged from 8-12cm when fully contracted up to 42cm when completely expanded (Figure 3A-B). To our perception, specimens had the brightest fluorescence but the dimmest bioluminescence among the 3 species. The light from *Veretillum* was green, though it appeared the most blue of the three species. Surprisingly, the fluorescence spectrum indicated the most red-shifted fluorescence (Figure 4) of the three species.

Green fluorescence was clear along the siphonozooids throughout the length of the rachis, as well as the tentacles of the autozooids (Figure 3C-F). Sites of fluorescence are also the same as those emitting light (Figure 3G, Supplemental Video), as noted by [Titschack, 1964]. Autozooids could emit light alone if touched. However, touching the rachis initiated luminescence across the body, appearing as waves extending from the site of stimulation that triggered both siphonozooids and autozooids to produce brief flashes of light. These waves of light could occur in either direction along the rachis. The waves of light appeared slower than those of *P. rubra*, and [Titschack, 1964] had estimated the speed at 14-15cm per second, though this depends on the contraction state of the colony.

## Discussion

### Role of GFP and green bioluminescence

All three species were dimly fluorescent under blue light, and required a filter to see the fluorescence (both by eye and with the camera). Nonetheless, the three spectra that were obtained indicate the presence of GFP. Based on our observations, there was complete correspondence between the fluorescence and bioluminescence in all three species, confirming the results by [Titschack, 1966]. This suggests that, like in *Renilla* [Morin and Hastings, 1971, Ward and Cormier, 1978], the GFP interacts closely with the luciferase. For many species, the native luciferases produce blue light [Wampler et al., 1973], which is converted to green light through the interactions with GFP. As it was theorized that the last common ancestor of cnidarians possessed a GFP [Shagin et al., 2004], this suggests that many species retained the ancestral GFP. As many sea pens are luminous and have high correspondance of fluorescence and bioluminescence, it appears that the GFP in sea pens is mostly used for energy transfer in bioluminescence. However, some species may have changes in the expression of their GFP gene (producing green or blue light), lost the GFP gene (producing blue light), or lost bioluminescence altogether.

Most sea pens inhabit soft substrate, though a few have been reported on rock [Williams and Alderslade, 2011]. Softer substrates may be habitats with higher sedimentation rates, or with strong currents that stir up sediment. While blue light is generally favored in the marine environment [Hastings, 1996], high amounts of particles may scatter light. Such scattering effect was suggested to have an influence on the evolution of the color of the light emission [Johnsen et al., 2012,Bessho-Uehara et al., 2020]. Across luminous Pennatulacea, 18 genera are described as luminous (Table 1), and most of these produce green light. An undescribed *Funiculina* species was reported to make blue light [Bessho-Uehara et al., 2020], while several species of *Umbellula* were reported to produce both blue and green light in a tissue-specific manner [Widder et al., 1983]. The species *Virgularia mirabilis* also was stated to produce blue light [Nicol, 1958], though no spectral data or photographs were reported. Thus, among luminous species, most produce green light.

**Table 1:**
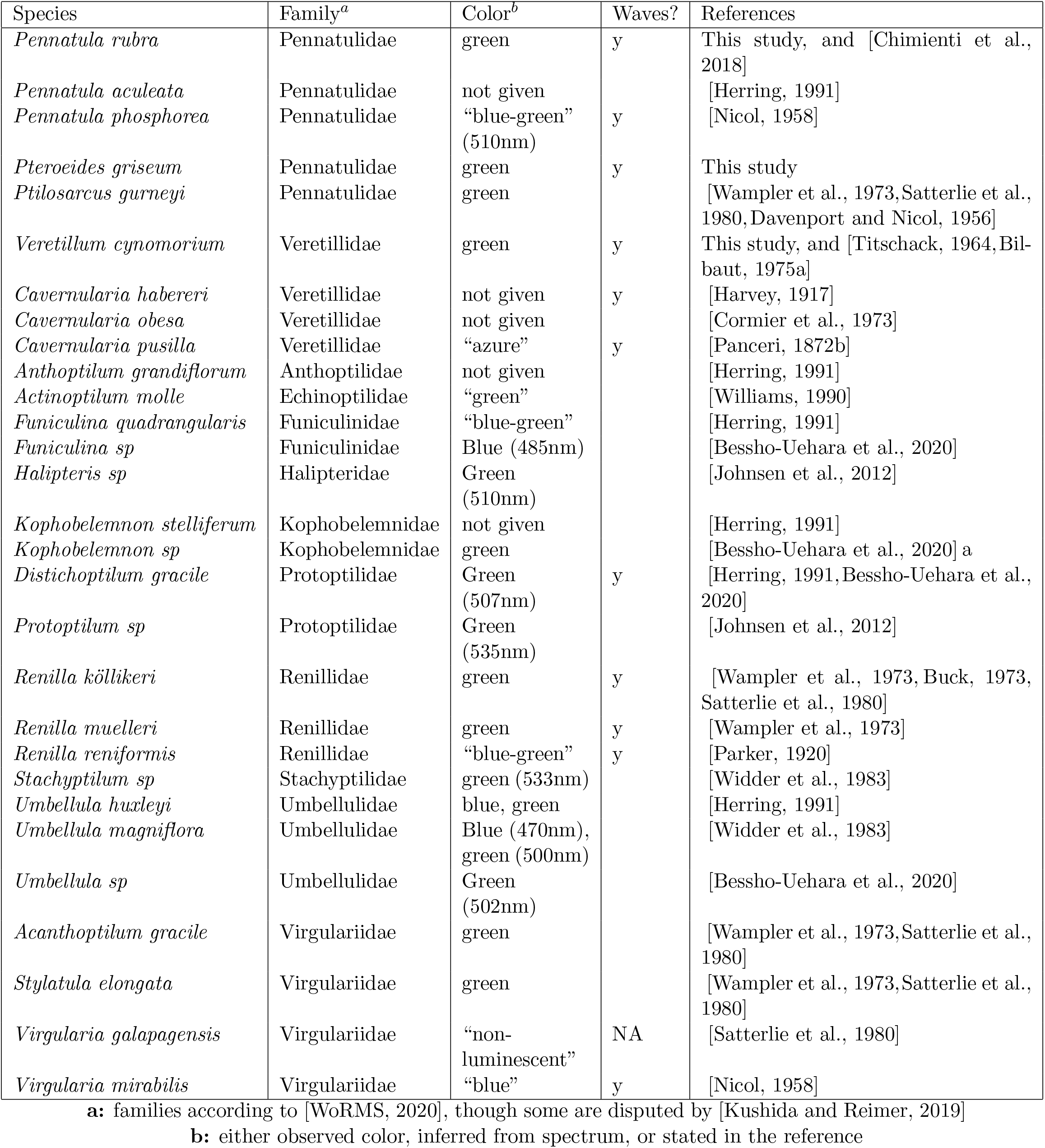
Summary of bioluminescence across Pennatulacea, expanded from [Williams, 2001]

| Species | Family<sup>a</sup> | Color<sup>b</sup> | Waves? | References |
| --- | --- | --- | --- | --- |
| *Pennatula rubra* | Pennatulidae | green | y | This study, and [Chimienti et al., 2018] |
| *Pennatula aculeata* | Pennatulidae | not given |  | [Herring, 1991] |
| *Pennatula phosphorea* | Pennatulidae | “blue-green”<br>(510nm) | y | [Nicol, 1958] |
| *Pteroeides griseum* | Pennatulidae | green | y | This study |
| *Ptilosarcus gurneyi* | Pennatulidae | green |  | [Wampler et al., 1973, Satterlie et al., 1980, Davenport and Nicol, 1956] |
| *Veretillum cynomorium* | Veretillidae | green | y | This study, and [Titschack, 1964, Bilbaut, 1975a] |
| *Cavernularia habereri* | Veretillidae | not given | y | [Harvey, 1917] |
| *Cavernularia obesa* | Veretillidae | not given |  | [Cormier et al., 1973] |
| *Cavernularia pusilla* | Veretillidae | “azure” | y | [Panceri, 1872b] |
| *Anthoptilum grandiflorum* | Anthoptilidae | not given |  | [Herring, 1991] |
| *Actinoptilum molle* | Echinoptilidae | “green” |  | [Williams, 1990] |
| *Funiculina quadrangularis* | Funiculinidae | “blue-green” |  | [Herring, 1991] |
| *Funiculina sp* | Funiculinidae | Blue (485nm) |  | [Bessho-Uehara et al., 2020] |
| *Halipteris sp* | Halipteridae | Green<br>(510nm) |  | [Johnsen et al., 2012] |
| *Kophobelemnon stelliferum* | Kophobelemnidae | not given |  | [Herring, 1991] |
| *Kophobelemnon sp* | Kophobelemnidae | green |  | [Bessho-Uehara et al., 2020] a |
| *Distichoptilum gracile* | Protoptilidae | Green<br>(507nm) | y | [Herring, 1991, Bessho-Uehara et al., 2020] |
| *Protoptilum sp* | Protoptilidae | Green<br>(535nm) |  | [Johnsen et al., 2012] |
| *Renilla köllikeri* | Renillidae | green | y | [Wampler et al., 1973, Buck, 1973, Satterlie et al., 1980] |
| *Renilla muelleri* | Renillidae | green | y | [Wampler et al., 1973] |
| *Renilla reniformis* | Renillidae | “blue-green” | y | [Parker, 1920] |
| *Stachyptilum sp* | Stachyptilidae | green (533nm) |  | [Widder et al., 1983] |
| *Umbellula huxleyi* | Umbellulidae | blue, green |  | [Herring, 1991] |
| *Umbellula magniflora* | Umbellulidae | Blue (470nm),<br>green (500nm) |  | [Widder et al., 1983] |
| *Umbellula sp* | Umbellulidae | Green<br>(502nm) |  | [Bessho-Uehara et al., 2020] |
| *Acanthoptilum gracile* | Virgulariidae | green |  | [Wampler et al., 1973, Satterlie et al., 1980] |
| *Stylatula elongata* | Virgulariidae | green |  | [Wampler et al., 1973, Satterlie et al., 1980] |
| *Virgularia galapagensis* | Virgulariidae | “non-luminescent” | NA | [Satterlie et al., 1980] |
| *Virgularia mirabilis* | Virgulariidae | “blue” | y | [Nicol, 1958] |

a: families according to [WoRMS, 2020], though some are disputed by [Kushida and Reimer, 2019]
b: either observed color, inferred from spectrum, or stated in the reference

Few species of Pennatulacea are specifically asserted to be non-luminous, including *Virgularia galapagensis* [Satterlie et al., 1980]. This is surprising because the related *V. mirabilis* is luminous. As [Nicol, 1958] had described much difficulty in reliably acquiring specimens of *V. mirabilis* that were in good shape to produce light, suggesting that they may be very sensitive to the disturbances of capture, or changes in temperature being brought to the surface. Thus, the luminous capacities of *V. galapagensis* may need to be re-examined. There are 19 remaining genera of Pennatulacea, which are *Amphiacme, Amphibelemnon, Calibelemnon, Cavernulina, Chunella, Crassophyllum, Echinoptilum, Gilibelemnon, Grasshoffia, Gyrophyllum, Lituaria, Malacobelemnon, Porcupinella, Ptilella, Sarcoptilus, Sclerobelemnon, Scleroptilum, Scytaliopsis*, and *Scytalium*. One of these (*Calibelemnon*) was incompletely identified, and is potentially luminous [Johnsen et al., 2012]. In fact, the remaining genera may be luminous as well, but appear to not have been examined.

### Potential function of bioluminescence in Pennatulacea

Many features of the bioluminescence of the pennatulacea in our study are strongly suggestive of a “burglar alarm” mechanism [Burkenroad, 1943]. This predicts that the light is produced when attacked by a first-order predator, and serves to signal for help to a second-order predator. Considering possible functions of the light in Pennatulacea, [Morin, 1976] listed several qualities that are displayed in the species in our study: (1) it is intermittent, i.e. not a continuous glow; (2) it is bright and conspicuous, i.e. showing a pattern of light, which varies by species; (3) it demonstrates an orientation or shape of the animal, i.e. overall the entire body emits some light to indicate its position and length; (4) it shows movement away from the stimulus, i.e. indicating precisely what site on the animal is being stimulated, or bitten; (5) it is proportional to the stimulus, i.e. can distinguish between being accidentally bumped versus being attacked. All three species in our study match these criteria.

Often following light production, colonies would contract into the aquarium sand, sometimes to the point of complete burial, a behavior also observed in the field [Chimienti et al., 2018]. Like light production, the contraction of the colony was faster with stronger physical stimulation. This contraction mechanism could take place directly after or in parallel to the burglar alarm. Assuming that a secondary predator does not arrive immediately, the sea pen could effectively hide or escape by contracting until it sinks into the sediment. However, in situ studies have shown that most specimens of *Ptilosarcus* would rapidly burrow into the sediment after subsequent physical contact with a predator [Weightman and Arsenault, 2002], rather than producing light. It is therefore possible that distinct conditions may favor one response over the other, perhaps by time of day or other cues from different predators.

Naturally, bioluminescence may also serve other functions in parallel, which are not mutually exclusive. Light production also could be an aposematic signal serving to ward away potential predators, indicating that the prey is toxic, unpalatable, or generally not worth continuing an attack. To us as observers, this may appear very similar to a burglar alarm. Given that light in sea pens is produced upon stimulation, not before, a predator would have already begun to disturb or try to eat the sea pen in order to see the light. Thus, some of the distinction may depend on the nature of the predator and prey.

An ecological survey on the luminous sea pen *Ptilosarcus gurneyi* had described seven common predators of *P. gurneyi* in the Puget Sound region of the northwestern United States [Birkeland, 1974]. Three of these predators (the seastars *Hippasteria spinosa* and *Dermasterias imbricata*, and the nudibranch *Armina californica*) were observed feeding only on *P. gurneyi*, while the other five were sometimes observed feeding on other things. For the specialist predators, bioluminescence of the sea pen could not be aposematic, as there would be no other food options to eat instead. In this case, it may serve to call a secondary predator, plausibly some nocturnal, visual predator, such as a crab. However, for the opportunistic predators attacking at night, light production could be aposematic and ward away these predators, as they have alternate food sources.

The nudibranch *A. californica* is also observed in southern California, and instead preys upon another Pennatulacean, *Renilla kollikeri* [Kastendiek, 1982]. The patterns of light emission are similar between the sea pens, as well as the wavelength. Because *R. kollikeri* also produces green light when disturbed, light again may function both as a burglar alarm for specialist predators, and as an aposematic signal for opportunistic predators. If most sea pens produce green light, then this may reinforce that color of bioluminescence as a warning signal, even if the predators can only see the color and intensity and not the pattern of light.

This same relationship occurs in the Mediterranean as well, as *Armina maculata* appears to feed mostly on *V. cynomorium* [Rosa et al., 2019]. Because *A. maculata* has a color and texture pattern that resembles a partially contracted *V. cynomorium*, then *A. maculata* would appear camouflaged when feeding on *V. cynomorium* during the day. However, it seems unlikely that the camouflage provides any defense at night, and so light production from the *V. cynomorium* may function here as a burglar alarm against *A. maculata*, among other possible predators.

### Distinguishing startling from a burglar alarm

It is difficult to separate a burglar alarm function from a first-order predator simply being scared by the light. Without knowing the mind of the predator, the observations may nonetheless be the same to us, i.e. that light is produced by the sea pen and the predator flees. In one case, the light itself is inherently aversive, while in the other case, the light indicates a more complex situation, whereby the position of the predator is advertised, and it flees to avoid being eaten itself. Even if a first-order predator were unaware that it was evading its own predator, evolution of a photophobic response may ultimately be due to selection by second-order predation. In this way, the burglar alarm is a transitory event, and is inherently difficult to observe. If this occurs at night, or in the deep sea, it may be even less likely to be observed by divers, or by ROVs, which will typically be operated during the day. Hence, this ultimately may require direct observations, perhaps by passive monitoring systems [Widder et al., 2005, Widder, 2007] set up in sea-pen fields.

## Conclusions

All three species were observed to produce “waves” of green light when disturbed, similar to many other sea pens around the world. As sea pens lacks eyes and are filter feeders, it is unlikely that light is used to attract prey or signal to conspecifics, so it is plausible that bioluminescence functions to deter predators or attract the attention of a second order predator. In situ observation at night may be necessary to resolve the possible functions of light of these animals.

## Special Thanks

WRF would like to thank K. Attard, SHD. Haddock, M. Bessho-Uehara, M. Kaiser, and the staff at the OOB Biodiversarium for helpful advice. WRF would also like to thank the divers at the OOB for careful collection of specimens. This work was supported by the European Union Horizon 2020 research and innovation programme (grant no. 730984) ASSEMBLE Plus 7th Call Transnational Access project “OctoLux” to WRF, and by a VILLUM Experiment grant (no. 00028022) to WRF. The authors declare no conflicting financial interest.

## Author Contributions

WRF designed experiments and acquired funding. WRF and ASdV did the experiments, analyzed the data, wrote and revised the paper.

## Data Availability

Raw fluorescence spectra can be downloaded at https://bitbucket.org/wrf/biolum-spectra/src/master/cnidaria/. The supplemental video of the bioluminescence can be found at https://bitbucket.org/wrf/biolum-spectra/downloads/ as either .mkv or .mp4.

